# Predicting mechanical ventilation effects on six human tissue transcriptomes

**DOI:** 10.1101/2022.01.19.476870

**Authors:** Judith Somekh, Nir Lotan, Ehud Sussman, Gur Arieh Yehuda

## Abstract

**Background:** Mechanical ventilation (MV) is a lifesaving therapy used for patients with respiratory failure. Nevertheless, MV is associated with numerous complications and increased mortality. The aim of this study is to define the effects of MV on gene expression of direct and peripheral human tissues.

**Methods:** Classification models were applied to Genotype-Tissue Expression Project (GTEx) gene expression data of six representative tissues– liver, adipose, skin, nerve-tibial, muscle and lung, for performance comparison and feature analysis. We utilized 18 prediction models using the Random Forest (RF), XGBoost (eXtreme Gradient Boosting) decision tree and ANN (Artificial Neural Network) methods to classify ventilation and non-ventilation samples and to compare their prediction performance for the six tissues. In the model comparison, the AUC (area under receiver operating curve), accuracy, precision, recall, and F1 score were used to evaluate the predictive performance of each model. We then conducted feature analysis per each tissue to detect MV marker genes followed by pathway enrichment analysis for these genes.

**Results:** XGBoost outperformed the other methods and predicted samples had undergone MV with an average accuracy for the six tissues of 0.951 and average AUC of 0.945. The feature analysis detected a combination of MV marker genes per each tested tissue, some common across several tissues. MV marker genes were mainly related to inflammation and fibrosis as well as cell development and movement regulation. The MV marker genes were significantly enriched in inflammatory and viral pathways.

**Conclusion:** The XGBoost method demonstrated clear enhanced performance and feature analysis compared to the other models. XGBoost was helpful in detecting the tissue-specific marker genes for identifying transcriptomic changes related to MV. Our results show that MV is associated with reduced development and movement in the tissues and higher inflammation and injury not only in direct tissues such as the lungs but also in peripheral tissues and thus should be carefully considered before being implemented.

## Introduction

Mechanical ventilation (MV) is a lifesaving intervention used for patients with respiratory failure. Although its therapeutic effects are well known, MV is also associated with numerous complications including significantly higher infection rates and lung injury, which prolong the duration of time spent on the ventilator and increase mortality [1], which can exceed 24% of those on ventilators [14]. The 2019 COVID-19 outbreak, resulting in MV treatment for numerous patients, has moved the question regarding invasive ventilation usage [22] to center stage.

### Clinical and transcriptomic effects of MV on tissues

The connection between treatment with invasive MV and pulmonary infections is so pervasive that the terms Ventilator-Associated Pneumonia (VAP) and ventilator-associated events (VAE) [2] were coined. VAE includes all the complications related to mechanical ventilation, broadening the horizon of possible consequences beyond those infection-related [2]–[4]. An example of VAE is the weaking of the diaphragm muscles [5], [6] that were observed after only twelve hours on an mechanical ventilator [5]. A large-scale study of 549 patients showed that blood samples of patients on MV exhibited increased lung inflammation and injury by testing inflammatory markers related to lung injury derived from blood samples [7]. MV contributes to mortality by inducing an inflammatory response in the lungs similar to that observed in acute respiratory distress syndrome (ARDS) [2], which can lead to multisystem organ failure. Although the most obvious clinical abnormalities in ALI (acute lung injury)/ARDS are related to the lungs, the most common cause of death is not due to hypoxia but to multiple organ dysfunction syndrome (MODS) [8]. Indeed, MV has been associated with greater risk of kidney failure [9], and diminished neurocognitive function in the brain [10]. Pinhu et al. [11] suggested two possible mechanisms through which MV induces multiple organ failure that are related to VAP and lung injuries that reduce the rate of organ perfusion.

Better understanding of the pathophysiology leading to the development of MODS in patients on MV should help in the development of approaches to interrupt the cascades leading to the syndrome [8]. The exact molecular mechanics of how MV affects peripheral organs is less obvious and the understanding of gene expression alteration in patients on MV may help. The ramifications of MV on human tissues’ gene expression are poorly explored and have focused mainly on the lungs [12]– [14]. MV was shown to stimulate the expression of the SARS-Cov-2 receptor ACE2 in the lungs [12] and a large-scale genomic research explored the effects of MV on the lung transcriptome [12]–[14]. The work in [13] used the GTEx gene expression data to detect several distinct gene expression clusters in the lungs, including a large cluster of genes associated with type II pneumocytes related to cells that proliferate in ventilator associated lung injury. Human and animal studies have demonstrated that MV using large tidal volumes (≥12 ml/kg) induces a potent inflammatory response and can cause acute lung injury [14]. Using a mouse model, [14] showed that non-injurious MV on its own initiates a proinflammatory transcriptional program in the lung. They compared breathing mice and mice on non-injurious MV (tidal volume of 10 ml/kg) and undertook an unbiased approach to partially decipher the complex network of related pathways. They showed that the low tidal volume still activates a transcription program of severe lung injury. In their previous work [15], they showed that the combination of non-injurious MV and low-dose exposure to bacterial products can cause severe lung injury in mice, implying a comodulatory role for MV in lungs that are at risk. MV was shown to affect gene expression of the lung areas [16]. While MV directly affects the lungs, investigating specific mechanistic effects on peripheral non-direct tissues has only been partially explored [8]–[10]. In addition, a large-scale investigation of MV-related changes in gene expression and molecular pathways in peripheral tissues is lacking.

### Machine learning methods for predicting MV and mortality

Machine learning models have been widely used in medical applications, including to predict MV and mortality. For example, several machine learning methods were used for predicting mortality of patients with acute kidney injury hospitalized in the ICU [17]. Artificial Neural Networks (ANNs or NNs) were used on data of breathing patterns to predict asynchronous breathing (AB) during MV [18]. Machine learning models used clinical data to predict MV mortality of COVID-19 patients in emergency rooms and in-hospital once the patient was admitted [19].

Machine learning approaches have been used not only to predict but also to detect the set of features that drive the prediction, e.g., analysis of gene expression data to discover novel marker genes, gene signatures and related pathways and networks, to differentiate between conditions. Machine learning can yield a list of differential genes that combine to drive the prediction and consider the dependence between the genes. For example, Grunwell et al. [22] applied machine learning to nanostring transcriptomics on primary airway cells and a neutrophil reporter assay to discover gene networks differentiating pediatric acute respiratory distress syndrome from non-pediatric ARDS. Cai et al. used gene expression data fed into three modeling methods—logistic regression, random forest and neural network—to develop a diagnostic gene signature for the diagnosis of VAP [23].

The Genotype-Tissue Expression (GTEx) [24] Project is a comprehensive public resource that includes tissue-specific gene expression data from nearly 1000 relatively healthy post-mortem and some material from “normal” surgical specimen human donors. The GTEx data of several tissues were successfully used in developing a machine learning model to predict the time since death of the donors [25].

In this study, we analyzed the GTEx [24] RNA-sequencing data to investigate the impact of MV on the transcriptomes of six representative human tissues. The study objective was to define gene signatures and pathways differentially expressed in direct and peripheral tissues of patients on MV. We constructed three machine learning models and analyzed the important features to predict patients who were on MV and those who were not, using the analysis of the transcriptomes to detect the significant MV-related gene signatures. Specifically, our models predict whether the donor was subject to ventilation prior to death. We trained and evaluated models on gene expression data of hundreds of samples for each tissue and thousands of genes as features.

To the best of our knowledge, ours is the first study to describe an investigation of the genomic effects of MV on peripheral tissues using machine learning algorithms and enrichment analysis of genes across tissues in human donors using gene expression data.

## Methods

### Data preprocessing

GTEx RNA-Seq data of 54 human tissues and 17382 RNA-seq samples from nearly 1000 donors was downloaded from the GTEx database (https://www.gtexportal.org/home/datasets, v8), and their transcript per million (TPM) values were log2-transformed. 18,680 protein-coding genes were retained. Outlier samples were filtered and all genes within each tissue were quantile normalized (to remove background and sample effects). For outlier removal, for each tissue we removed outlier samples by applying an Isolation Forest algorithm [26], with a parameter of 0.01. Accordingly, the 1% of the remaining samples that had the highest levels of variation from the other samples were also removed. Genes with zero variance were excluded from the calculation (e.g., for adipose– subcutaneous, 262 genes were excluded). For a given tissue, genes having at least 0.1 TPM in 80% or more of the samples were retained. Supplemental Table S1 presents the number of samples and features per tested tissue type after preprocessing and correcting for confounding factors. For example, for adipose–subcutaneous, there are 544 samples; for each sample, there are 16,052 genes that we use as features in the machine learning models.

### Confounding factor adjustment

Our previous work [27] showed that linear regression-based adjustment of the heterogenous GTEx data outperforms other methods in preserving the biological signal—which is relevant here. Thus, we used linear regression models to adjust for the known confounding factors—experimental batch, ischemic time (time elapsed between actual death and sample extraction), gender and age. Age covers the 20–80-year range and is partitioned into 10-year intervals (embedded in the GTEx dataset).

The type of death classification of the samples (DTHHRDY=death circumstances) is based on a four-point Hardy Scale. 0 represents cases on mechanical ventilator prior to death, 1 and 2 represent non-ventilation deaths (short and intermediate duration prior to death) of healthy individuals, and 3 and 4 represent non-healthy individuals. As the focus of the research is on healthy individuals at the time of death, we excluded samples with a DTHHRDY value of 3 “Intermediate death for ill patients” and 4 “Slow death” that represent non-healthy individuals with a long-term illness (these also comprised a small number of samples).

We aggregated the DTHHRDY into two categories—death type 0 (ventilation), a subject who was on a ventilation machine prior to death and death type 1 (non-ventilation), a subject who was not connected to a ventilator at the time of death. Ischemic time is the time in minutes that elapsed between death and sample extraction. We found that there was a correlation between the ischemic time (SMTSISCH) and DTHHRDY (ventilation vs. non-ventilation) that wrongly skewed the ischemic time coefficient when we used both as predictors in the linear regression model (see explanations and plots in Figures S3 and S4 in supplemental file S1). This phenomenon is the result of the fact that individuals on MV were already in the hospital and this resulted in shorter time (ischemic time) between time of death and sample collection. This phenomenon was detected previously [25] and dealt with by correcting for only one of these two confounding factors. Here we developed an improved approach to correct the data for ischemic time but with minimal harm to the ventilation signal. We performed linear regression in two steps. We first corrected for age, sex and batch and then for ischemic time, by inferring its coefficient for each group separately; thus, we did not skew by the correlated death type. We used the averaged coefficients calculated for each group (ventilation/non-ventilation) independently as explained below. After correcting the data with our linear regression model, we used the residuals as the expression values for further analysis.

The two-step process of extracting residuals was as follows:

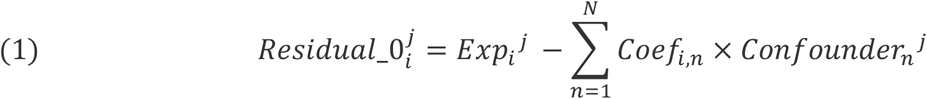

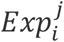 is the expression level of gene *i* in sample *j. Coef*_*i,n*_ is the multiple linear regression coefficient gene *i* in coefficient *n*. 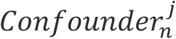 is the phenotype confounding value for sample j and confounding factor n. 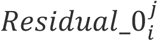 is the residual level of the value for gene i and sample j. The confounding factors in the model and their corresponding confounding coefficients were gender, age, and the experimental batch. Then, the ischemic rate coefficient was calculated for each two ventilation types separately in order to correct independently for ischemic time and not for ventilation type (which are correlated). We separated the residuals by the two ventilation types (ventilation and non-ventilation). An unweighted average of the coefficients by ventilation type *Ischem*_*i*_ per gene, was taken. The residual used was:

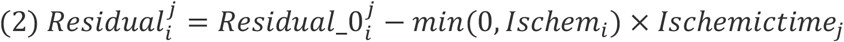

*Ischem*_*i*_ is the average of the factor calculated for ischemic time for death type 0 (ventilator) and death type 1 (non-ventilator). If the average of the ischemic coefficients was greater than 0, then there was no further adjustment performed for ischemic time. *Ischemictime*_*j*_ is the ischemic time for sample j. 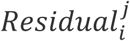 is the residual value once the two-step regression process was completed for gene i and sample j. Ideally, we would have liked to perform the regression in one step; however, we found that for the genes with the highest predictive rates for ventilation type, this would have yielded skewed the ischemic time coefficients. Further details and explanatory plots of the correction approaches we tested and the final two-step linear regression approach we used can be found in supplemental file S1.

### Machine learning methods

We selected three different algorithms and six main tissues and built machine learning classifiers for each tissue—resulting in a total of 18 machine learning models. Each model is a binary classifier designed to predict the ventilation/non-ventilation samples based on the RNA-seq gene levels per samples, i.e., the features, derived from the specific tissue. The selection of algorithms was somewhat limited given the nature of the data: since ours was a relatively small sample size (∼200–700 items) having a large number of features (∼13-16k), not all machine learning methods were expected to achieve good results. Due to the relatively small sample size, we decided to experiment with two different decision tree-based algorithms: Random Forest (RF), and a boosted decision tree, the eXtreme Gradient Boosting technique (XGBoost) [28]. We expected that the boosted decision trees would be very effective in our context. Tree boosting machines are a family of powerful machine learning techniques that have shown considerable success in a wide range of practical applications. An additional advantage of tree boosting machines is the explainability capabilities of these models, which can help in validating the correctness of the model, by checking the relevance of the most significant gene levels to the tested conditions, i.e., ventilation vs. non-ventilation, and learning the biological signs that the model is detecting. In addition, and given the popularity of using Deep Learning and the extensive usage of it in many different use cases these days, we also chose to use the Artificial Neural Network (ANN) [29] as one of our methods, although it may be less effective when working with tabular data and a relatively small sample size [30].

### Random Forest

RF is an ensemble algorithm that combines multiple decorrelated decision tree prediction variables based on each subset of data samples [31]. RF has been extremely successful as a general-purpose classification and regression method, proven to be a computationally efficient technique that can operate quickly over large datasets, and is easy to implement. RF can also handle a large number of input variables without overfitting [32]. In general, the RF approach creates several randomized decision trees, then combines and aggregates their predictions by averaging.

The RF model was built using the scikit-learn^1^ Random Forest Regressor. We used a model configuration similar to the XGBoost (presented below), i.e., we use 100 estimators with a maximum depth of 3 and a learning rate of 0.1.

### Tree Boosting – XGBoost

*Boosting* [33] is a commonly used machine learning method that attempts to improve the accuracy of a given learning algorithm. Boosting is done by creating an ensemble learner from several weaker models. The ensemble takes a set of predictors, all aiming to predict the same target, and combines them together into to form a stronger predictor. Friedman [34] was the first to propose a gradient-descent-based formulation of boosting, a method that improves the approximation accuracy.

The XGBoost technique [28] method is based on Friedman’s gradient boosting but introduces additional improvements that increase the technique’s performance and the accuracy of its results. While in the original gradient boosting model, the trees are built in series, XGBoost does this in a parallel way, similar to the RF method that grows trees in parallel to each other, and each tree tries to compensate for the areas in which the previous tree was less accurate. This method also uses regularization terms to control the variance of the fit and control the flexibility of the learning task, while obtaining models that generalize better to unseen data. XGBoost has been extensively used recently and shown useful in solving different real-world problems, for example, pathway analysis of biomedical data [35] and diagnosis of chronic kidney disease [36].

We executed the XGBoost algorithm to create a set of 100 decision trees for each tissue. An example of one generated XGBoost tree and its branches for muscle-skeletal tissue classification is provided in supplemental Figure S1. Similarly, supplemental Figure S2 provides one of the tree branches that were generated for the adipose-subcutaneous tissue. After the tree is created, it can be used for prediction when each tree node, depicted by an ellipse as seen in supplemental Figures S1 and S2, that represents a condition on a gene expression value is checked for each predicted sample. For each tree node from the top of the tree, if the sample’s value equals the tree node specification, the selected path in the decision tree is the ‘yes’ path. Else the ‘no’ path is selected. If the value is missing, the ‘missing’ path is selected. For example, for the provided “Adipose–Subcutaneous” tree in supplemental Figure S1, the level of MXRA5 in the sample is compared to 0.32043466. If the value is smaller than 0.32043466, or there is no measurement, the left path is selected. Eventually the selected sample receives a score (for being ventilation/non-ventilation) for each tree. The scores of 100 trees will be combined to determine the selected ventilation/non-ventilation class. Each of the 100 trees may contain different genes and check different values for these gene levels.

To develop the XGBoost binary classifier we used the XGBoost Python library [37]. To avoid overfitting, while maintaining high performance predictions, we configured the XGBoost to use 100 estimators with a maximum depth of 3. We found that a high number of estimators (100) performed better than lower values (e.g., 30 or 50). A learning rate of 0.1 was found to be effective in this case (lower values were tested). For the rest of the parameters, the default values were used.

### Artificial Neural Network (ANN)

ANN [29] is a computational model inspired by biological models, which often exceed the performance of previous forms of artificial intelligence used in many common machine learning tasks. It is basically a data processing system composed of a combination of simple, interconnected processing elements, in a predesigned architecture. The ANN basic building block is a simple mathematical function that includes three steps: multiplication, summation and activation. For the purpose of this research, we created a network with a shallow topology of one hidden layer, as shown in supplemental Figure S6. Additional topologies were tested, but we found that a single hidden layer network provides the best results. The selected topology includes an input layer with the dimensions of the number of features (based on the relevant organ), a hidden layer of 100 nodes with a Rectified Linear Unit (ReLU) activation function [38], and a single output layer with a sigmoid activation function. We use binary cross-entropy as the loss function, and an adaptive moment estimation optimizer (Adam) [29].

The neural network model was designed using the Keras^2^ network, with TensorFlow^3^ as its backend. To confirm the effectiveness of the XGBoost model in predicting MV, we used ANN and RF, widely used machine learning models, for comparison and summarized the advantages and disadvantages of each model in Table 1.

**Table 1.** Comparison between the machine learning models used in the study.

| Model | Advantages | Disadvantages |
| --- | --- | --- |
| XGBoost | • Effective for a relatively small number of samples with a large number of features<br>• Encapsulated explainability capabilities that can help validate the correctness of the model, e.g., by checking the relevance of the most significant gene levels to the tested condition<br>• Includes improvements to the original gradient boosting model that increase the performance and the accuracy of the results | • May exhibit overfitting if hyperparameters are not adjusted correctly<br>• Applicable for numeric features only |
| RF | • Easy to implement both for classification and regression tasks<br>• Provides some level of explainability<br>• Avoids overfitting | • Lower performance than more modern methods<br>• Nonoptimal performance when classes are unbalanced |
| ANN | • Excels at cognitive tasks (image/video/text/voice data) | • Requires a large number of samples<br>• Hard to explain and detect the feature importance<br>• May be less effective with tabular data |

### Computation Resources and Frameworks

The training of the models was conducted on an Intel(R) Core(TM) i9-7920X CPU @ 2.90GHz computer with 24 CPUs, with 128GB RAM and an NVIDIA Corporation GV100 [TITAN V] (rev a1) GPU. Training in this setup lasted about five days.

### 10-fold cross-validation

For model performance evaluations, i.e., to evaluate whether the model is accurate and not overfitted and due to the relatively low number of samples (∼200–700 samples), we used K-Fold Cross Validation—more precisely, the scikit-learn^4^ implementation of Stratified K-Fold Cross Validation [39] without shuffling. This cross-validation is a variation of K-Fold that returns stratified folds. The folds are made by preserving the percentage of samples for each class. We chose a split of 10, so each time 90% of the data is used for training, and 10% for validation. We used the same fold and data when building the ANN, RF and XGBoost models to assure we compared the models under exactly the same terms.

For each organ we configured each model to use the organ’s gene levels as features, with the ventilation/non-ventilation types as the predicted target. We performed 18 experiments, each per distinct model and tissue. Each experiment included a 10-fold cross-validation prediction, and we saved the results of the 10 executions. We averaged the results of the 10 executions as well as the feature importance. For each prediction we tracked the average and standard deviation of the Area Under the ROC curve (AUC), accuracy, F1 score, recall and precision of each experiment.

### Feature analysis

Feature analysis was performed using Lundberg’s approach for explaining boosted trees called SHAP – SHapley Additive exPlanations [40]. SHAP offers a high-speed precise algorithm that can explain the output of any machine learning model, in particular tree ensemble methods. SHAP calculates values for each feature, representing how much each feature contributes to push the model output from the base value (the average model output over the training dataset we processed) to the model output.

### Pathway enrichment analysis

We imported a list of the top genes (feature score > 0.01) into the web-based Enricher tool [41] using KEGG 2021 pathways.

## Results

We used large-scale RNA-seq gene expression data from the GTEx project [42] for six representative human tissues—adipose-subcutaneous, liver, lung, muscle-skeletal, nerve-tibial and skin-sun exposed (lower leg) (see supplemental Table S1 for the full number of samples and genes for each tissue). We aggregated the samples into ventilation and non-ventilation groups as described in the Methods section. The tissues’ gene expression data was preprocessed and corrected for confounding factors as described in the Methods section and in the supplemental file.

To detect a combination of marker genes signifying MV usage, we created 18 prediction models using the RF, XGBoost decision tree and ANN methods to classify the ventilation and non-ventilation samples for six representative tissues. To evaluate the predictive performance of each model, we compared the classifiers using the AUC, accuracy, precision, recall and F1 scores. We finally conducted feature analysis per each tissue to detect MV marker genes, followed by pathway enrichment analysis for these genes.

### Classification comparison

Table 2 provides a detailed comparison between the methods performance in means of the different metrics: AUC, Accuracy, F1 Score, Recall and Precision. The numbers provided here are the average of the 10-fold executions across the six tissues. Detailed results including standard deviation between executions can be found in supplemental file Table S2. It is easy to see here and in Figure 1 that XGBoost outperforms the ANN and RF models in all metrics aside from recall, in which the ANN model outperforms XGBoost but only by a small margin. We added the AUC metric in our analyses since the class distribution within the data is proportional but not fully balanced (see Table S1 in the supplemental file for the number of samples in each class).

**Table 2:** Binary classification model evaluations.

| Model \ Evaluation | Average AUC | Average Accuracy | Average F1_score | Average Recall | Average Precision |
| --- | --- | --- | --- | --- | --- |
| Neural Net | 0.931 | 0.934 | 0.916 | **0.931** | 0.903 |
| Random Forest | 0.901 | 0.912 | 0.880 | 0.862 | 0.905 |
| XGBoost | **0.945** | **0.951** | **0.936** | 0.924 | **0.951** |

**Figure 1.**
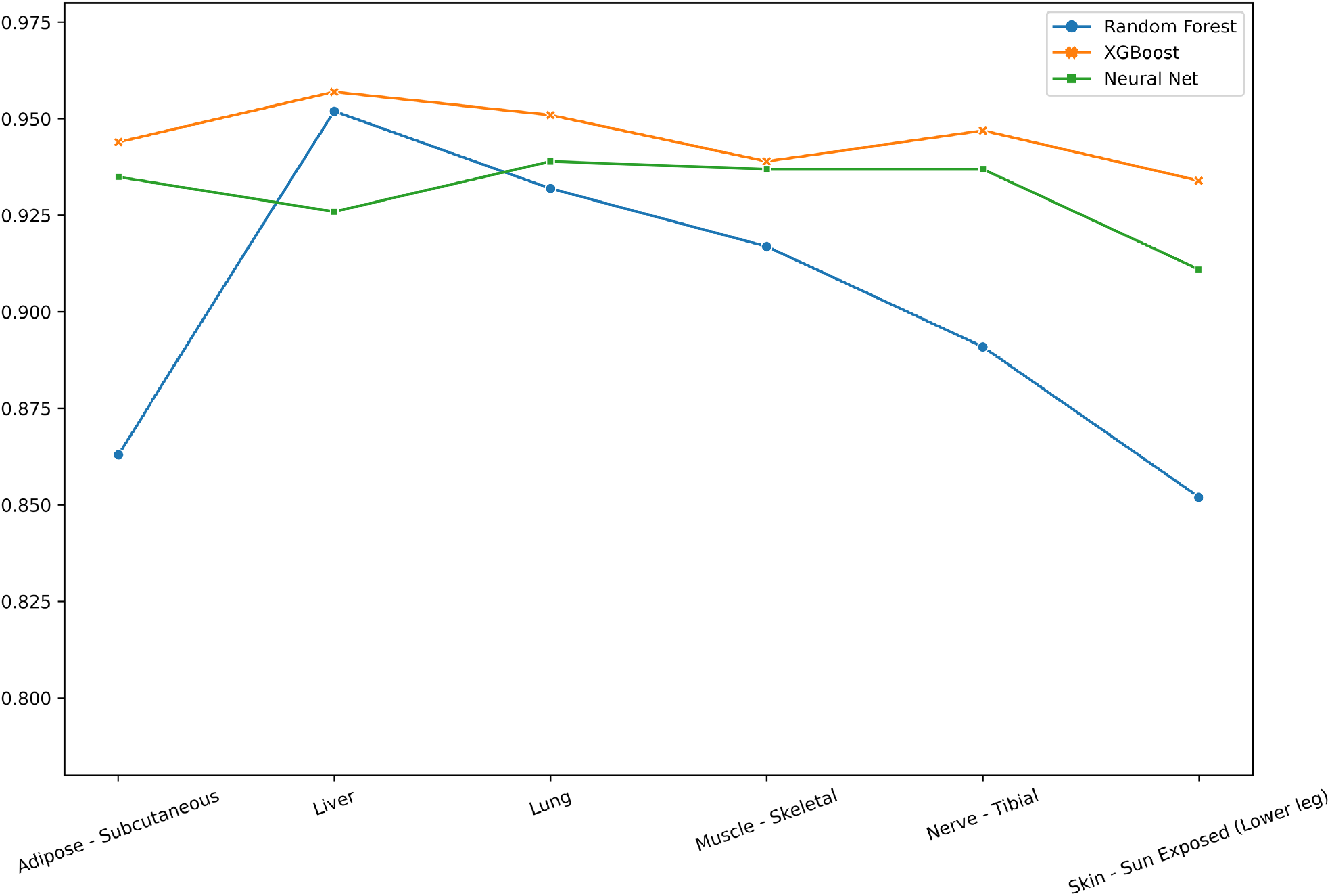
AUC comparison of the 18 classifiers. It can be seen that the XGBoost model outperforms the RF and ANN models for the six tested tissues.

Table 3 provides a comparison of the results of the 18 models’ performance, focusing on the AUC and providing the average AUC and AUC per organ. Clearly, even when looking at each tissue individually, XGBoost outperforms ANN and RF, although in some cases the difference is not significant. Given that XGBoost outperforms the two other methods (see also Figure 1), while providing good feature analysis capabilities, from this point on we focus on the modeling, experiment and results of the XGBoost model.

**Table 3:**
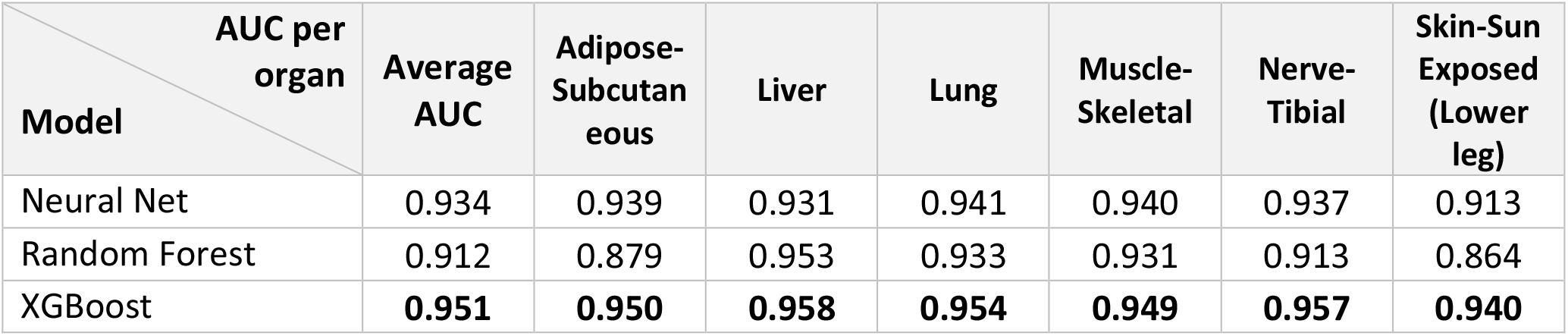
Binary classification models’ AUC for the different organs.

### Tissue-specific MV marker genes

To detect the most predictive and ventilation discriminant genes, for each tissue we performed feature analysis using SHAP [40] to explain the boosted trees. SHAP calculates values for each feature, representing how much each feature contributes to elevate the base value of the model, the average model output over the training dataset we processed, to the model output. Figure 2 demonstrates the feature analysis results for muscle-skeletal and lung and presents the mean absolute value of the SHAP values for each feature in our model. The top 20 features’ average SHAP impact on model output magnitude in absolute values are presented. A summary of SHAP scores for the top 20 genes in all tested tissues are provided in Table 4.

**Table 4.** Top 20 genes with highest importance SHAP scores in each tissue. Genes that are common to more than two tissues are highlighted in bold.

|  | Adi-<br>pose | Score | Liver | Score | Lung | Score | Muscle | Score | Nerve<br>Tibial | Score | Skin | Score |
| --- | --- | --- | --- | --- | --- | --- | --- | --- | --- | --- | --- | --- |
| 1 | **MXRA5** | 0.49 | MGMT | 0.63 | PHF13 | 0.30 | **GKN1** | 0.52 | FDCSP | 0.54 | AVP | 0.53 |
| 2 | VENTX | 0.42 | DNAJB<br>4 | 0.55 | MT4 | 0.29 | **CLEC3B** | 0.34 | HSD11<br>B2 | 0.50 | **TFF1** | 0.41 |
| 3 | **EGR1** | 0.31 | C5orf2<br>4 | 0.26 | HRG | 0.23 | TET1 | 0.26 | CRABP<br>1 | 0.44 | ODF3L<br>1 | 0.24 |
| 4 | PELO | 0.27 | TOR1A | 0.24 | TBC1D<br>22B | 0.23 | FGF6 | 0.21 | **EGR1** | 0.31 | **MXRA5** | 0.23 |
| 5 | **CLEC3B** | 0.25 | GATAD<br>1 | 0.22 | LCE5A | 0.20 | SMCO1 | 0.19 | SLN | 0.22 | **KRT1** | 0.23 |
| 6 | **KRT6C** | 0.22 | C12orf<br>60 | 0.14 | ALPK3 | 0.18 | SPSB4 | 0.18 | SLITRK<br>6 | 0.18 | **CYS1** | 0.20 |
| 7 | **CYS1** | 0.20 | GPRIN1 | 0.13 | **MXRA5** | 0.17 | ZCCHC<br>24 | 0.17 | **KRT20** | 0.18 | NEURL<br>2 | 0.19 |
| 8 | MUC21 | 0.20 | KCNJ8 | 0.13 | LCE2C | 0.16 | KRT6B | 0.17 | XIRP1 | 0.15 | RP11-676J12.7 | 0.19 |
| 9 | C22orf31 | 0.19 | DRD4 | 0.12 | CYP1A2 | 0.14 | CCND1 | 0.16 | HBQ1 | 0.14 | KRTAP5-6 | 0.17 |
| 10 | CX3CL1 | 0.17 | ENTPD7 | 0.12 | ADRA1B | 0.14 | EGR1 | 0.15 | BHLHE40 | 0.13 | NIPSNAP3A | 0.15 |
| 11 | C10orf99 | 0.15 | SLC25A21-AS1 | 0.11 | TERF2IP | 0.12 | TFF1 | 0.14 | SFRP2 | 0.13 | FLG | 0.13 |
| 12 | DEFA6 | 0.15 | OPN1SW | 0.10 | LCE2A | 0.12 | FITM1 | 0.13 | GUCA2A | 0.12 | RD3 | 0.11 |
| 13 | PRND | 0.14 | PAQR8 | 0.09 | SST | 0.11 | MRPL16 | 0.13 | CD248 | 0.12 | EGR3 | 0.10 |
| 14 | SMCP | 0.13 | STX11 | 0.09 | B3GNT2 | 0.10 | TLL2 | 0.12 | RP11-10J21.3 | 0.12 | ATP4B | 0.10 |
| 15 | TNFRSF21 | 0.09 | LRRC40 | 0.08 | DYRK2 | 0.10 | SPRR2A | 0.12 | DKK4.00 | 0.09 | WFDC12 | 0.10 |
| 16 | GKN1 | 0.09 | CECR6 | 0.08 | DPRX | 0.10 | CEACAM6 | 0.12 | RP11-268J15.5 | 0.09 | OCM2 | 0.10 |
| 17 | HUS1B | 0.09 | OMG | 0.08 | CHP2 | 0.09 | SLN | 0.11 | DUPD1 | 0.09 | DUSP1 | 0.10 |
| 18 | TIFAB | 0.09 | IFITM10 | 0.08 | AGTR2 | 0.09 | SPATA25 | 0.10 | TAS2R46 | 0.08 | SCNN1G | 0.09 |
| 19 | P2RY13 | 0.08 | TMEM200C | 0.07 | SPINT3 | 0.09 | SOCS4 | 0.09 | DBX2 | 0.08 | LGALS7B | 0.09 |
| 20 | TCF21 | 0.08 | PSPN | 0.07 | LSM11 | 0.09 | TBC1D12 | 0.09 | PSD2 | 0.08 | IER2 | 0.09 |

**Figure 2.**
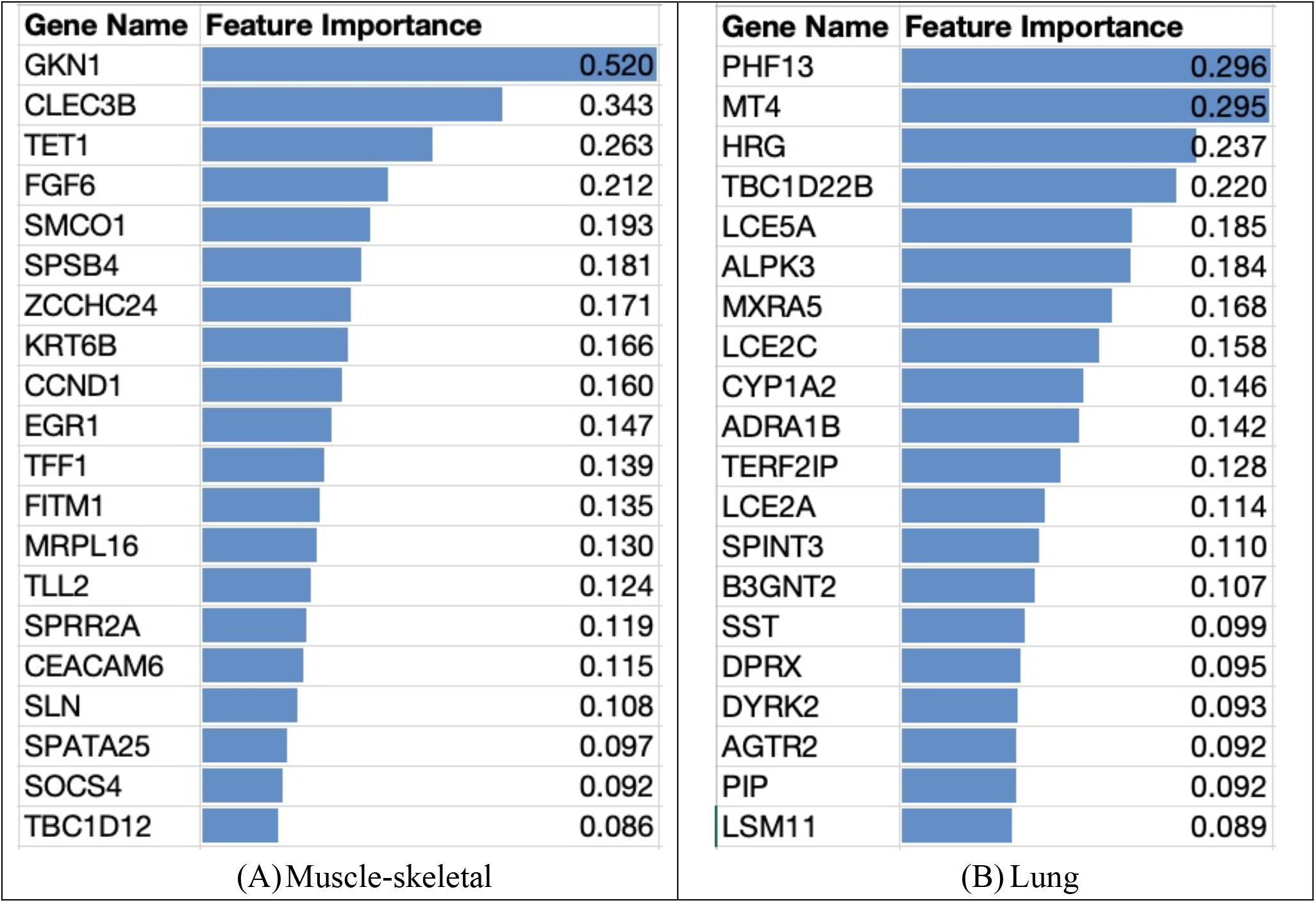
Top 20 genes and average SHAP impact (absolute values) on the magnitude of model classification output. (A) Values for muscle-skeletal tissue. It can be seen that GKN1 gene expression values have the highest impact on the MV classification. (B) Values for the lung tissue.

The SHAP approach can also explain how each feature (gene) contributes to the classification per class.

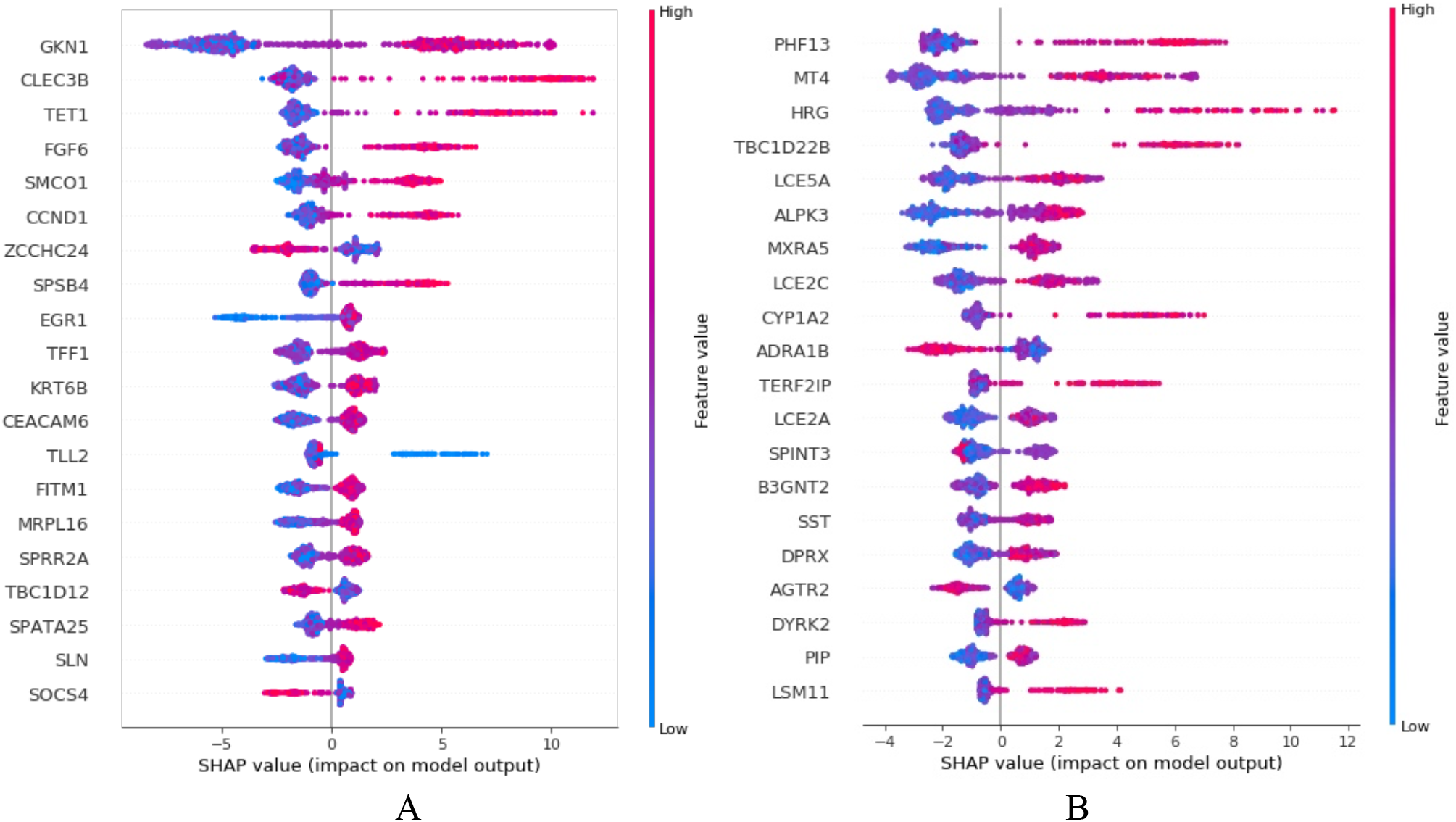

Figure 3 provides a plot of genes sorted in descending order by gene importance, representing the sum of SHAP value magnitudes over all samples and uses SHAP values to show the distribution of feature impacts on the model output. The horizontal location shows the impact of each feature, i.e., whether the effect of that value is associated with a higher or lower prediction. The colour relates to the original values of each gene across samples and shows whether that variable is high (in red) or low (in blue) for that observation. Red represents a higher value of the gene for the ventilation samples compared to the average values across all samples in the measured tissue; blue represents a low measured value. The x-axis in

**Figure 3.**
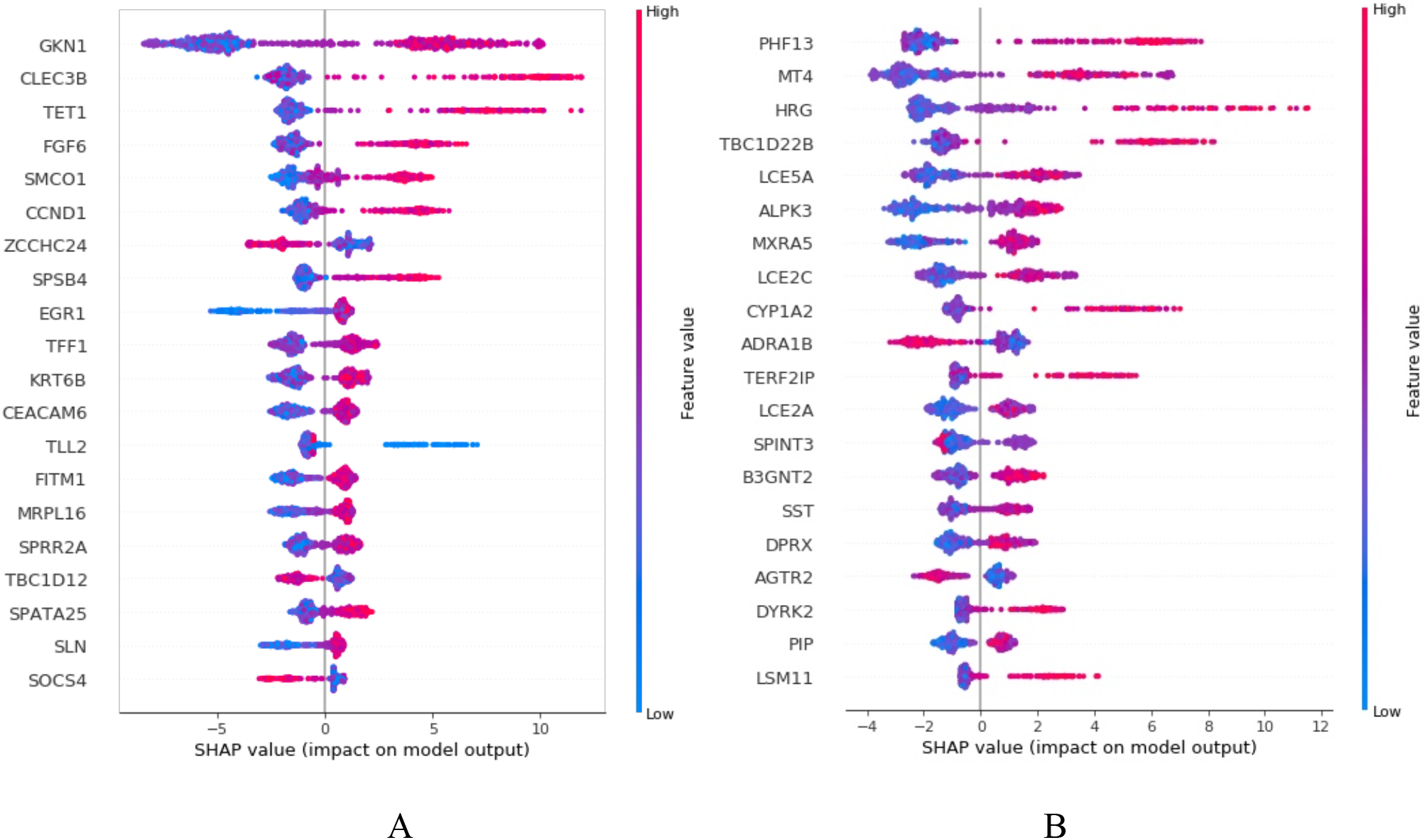
SHAP variable importance plots. (A) SHAP values for muscle-skeletal tissue. (B) SHAP values for the lung tissue. The plot includes all samples in the training data and the values represent the impact of the gene on model prediction output. SHAP values explain to what extent the feature (gene) contributes to the prediction of the model.

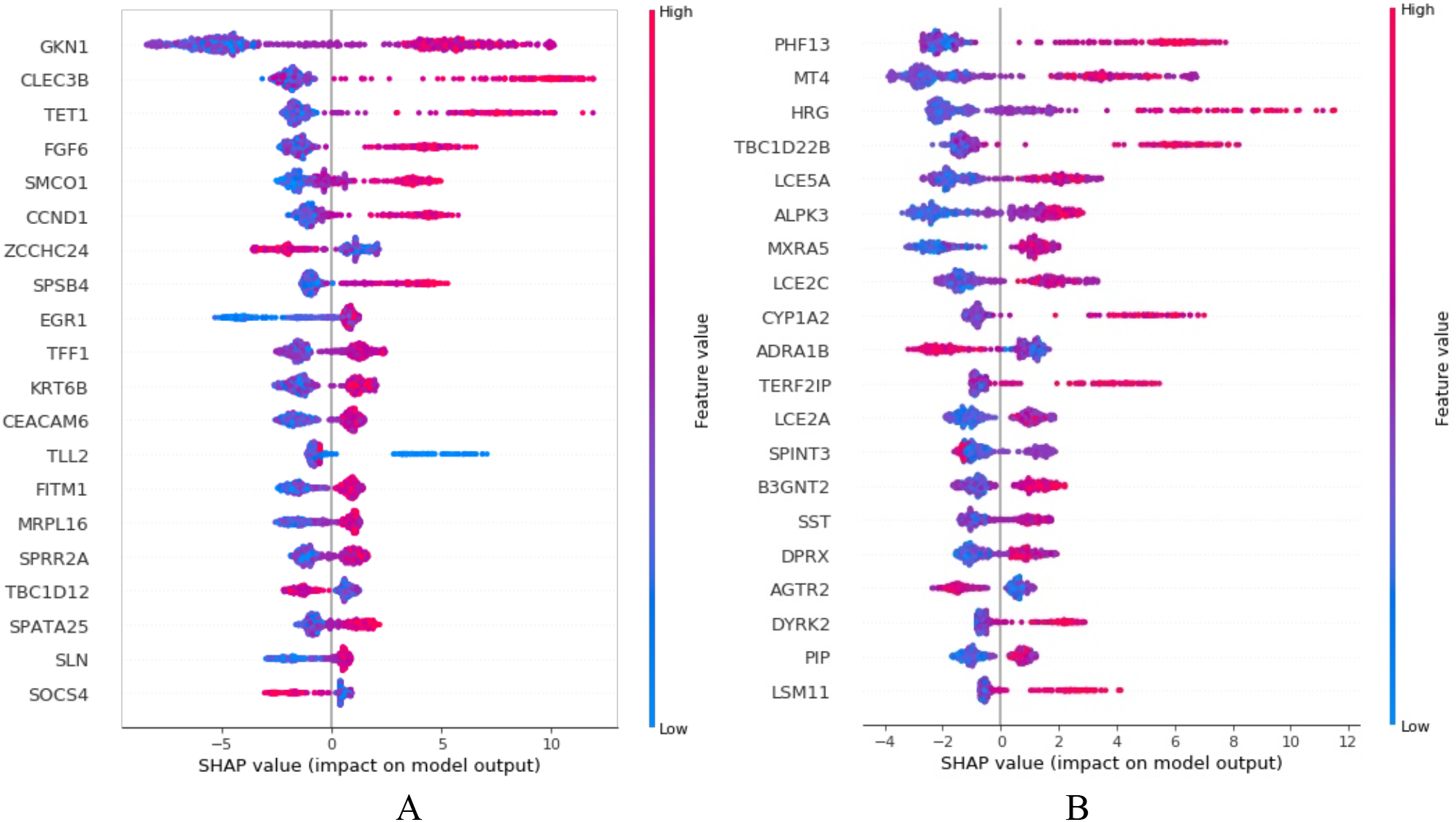

Figure 3 is the SHAP values of each gene and represents the impact of the gene on model output in muscle-skeletal and lung tissues. A low SHAP value means that this sample is more likely to be a ventilation sample and a high SHAP value means that this sample is more likely to be a non-ventilation sample. This approach reveals, for example, that for most cases, a high value of CLEC3B (the second gene from the top in

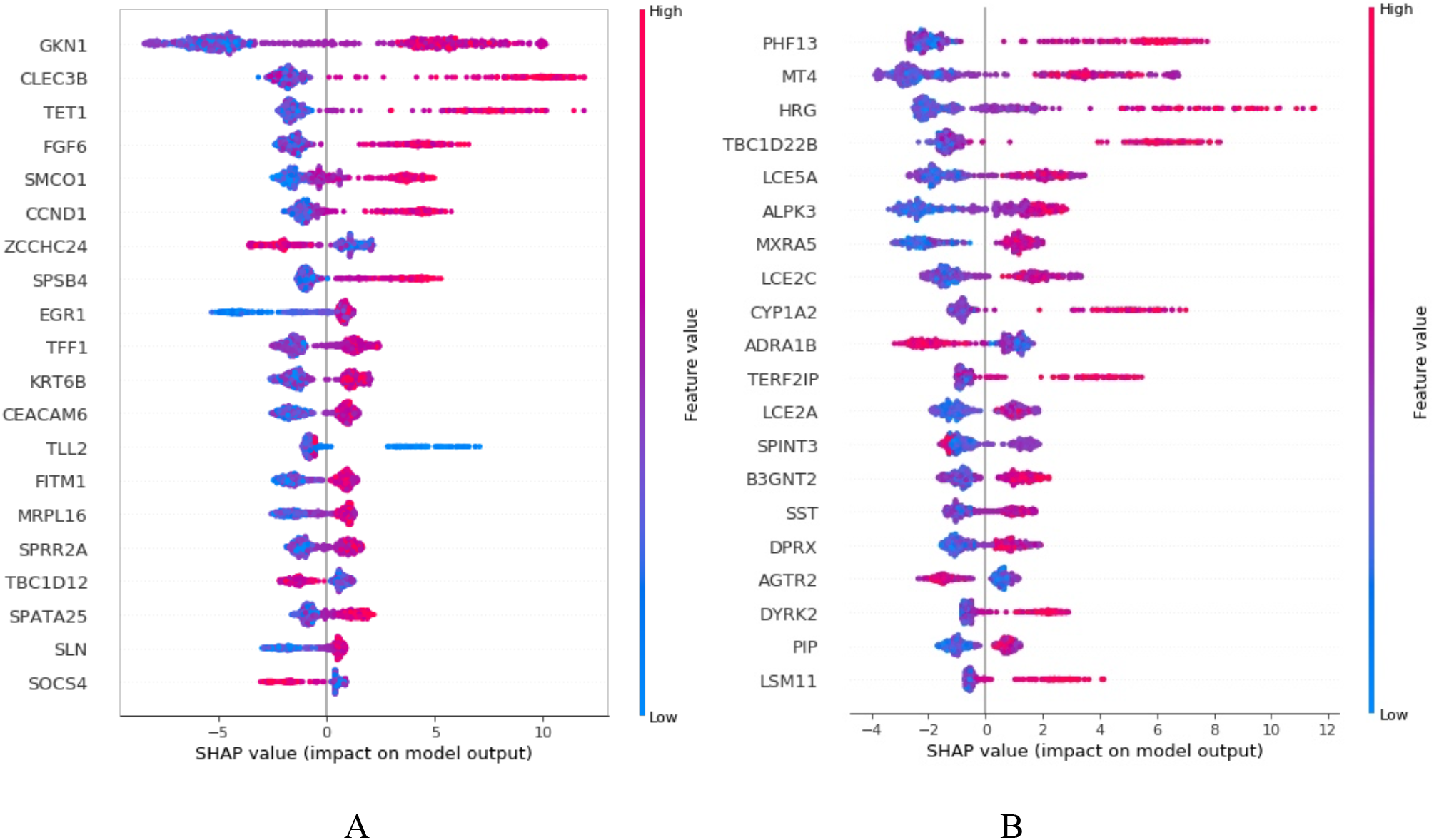

Figure 3A) has a high and positive impact on non-ventilation prediction and increases the chances of the sample to be classified as taken from a non-ventilator group. The high comes from the red color and the positive impact shown on the x-axis. The CLEC3B gene has been reported to regulate muscle development [43]. An example of negative correlation is the AGTR2 gene (fourth from the bottom in

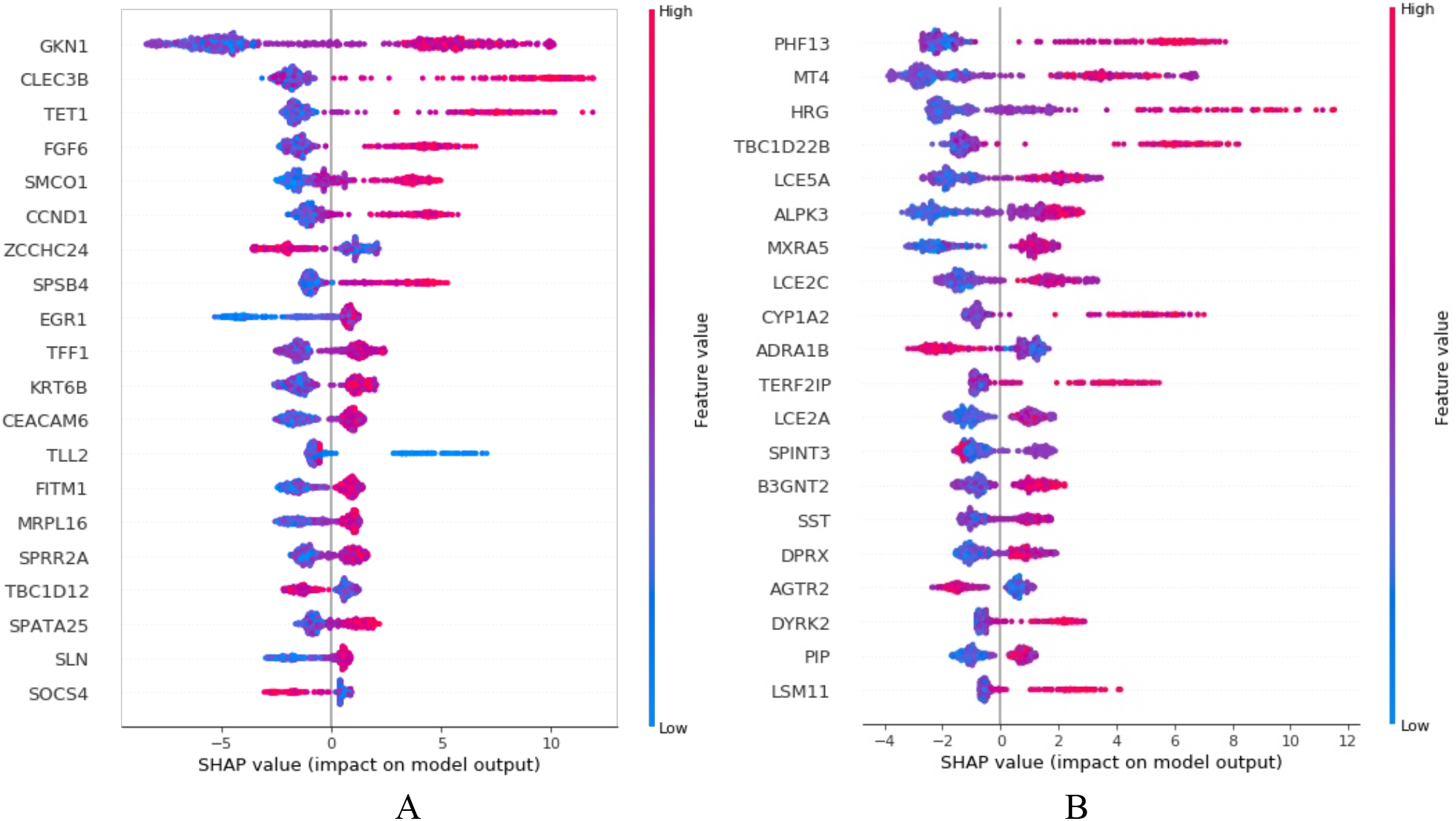

Figure 3B) that has a high and negative impact on the non-ventilation prediction, i.e., it is lower in the non-ventilation group and higher in the ventilation group. We note that the angiotensin II receptor 2 (AGTR2) gene [44] is expressed in lung fibrosis. The diagram also illustrates the importance of using multiple features and feature combinations for achieving the high accuracy. We note that the tree-based method is based on calculating threshold values for a combination of genes to drive the prediction. An example of such combination, an XGBoost tree branch, is presented in supplemental Figures S1 and S2 and explained in the Methods section. One feature is not enough to differentiate between the samples and a combination of features is required. It is easy to see, for example, that for muscle-skeletal, high values of GKN1 correlate in most cases to the non-ventilator group. This gene’s levels, however, are not good enough to be used as a single discriminator, since there are several cases of samples in the ventilator group that have high levels of GKN1. Only by using the combination of gene levels can the model achieve high accuracy/AUC. Additional histograms illustrating the differences in CLEC3B and AGTR2 gene values in muscle-skeletal and lung tissues for the ventilation and non-ventilation types are included in supplemental figure S3.

Table 4 summarizes the top 20 most predictive genes for the six tissues. Genes common to several tissues appear in bold.

We used the web-based Enricher tool [41] for pathway enrichment analysis using KEGG 2021 pathways of the top genes per tissue (feature score > 0.01). For example, we tested the enrichment of 149 genes for muscle-skeletal and 148 genes for adipose-subcutaneous. The “Cytokine-cytokine receptor interaction” pathway was significantly enriched (adjusted p-value < 0.05) in four tissues— adipose-subcutaneous, liver, muscle-skeletal and skin. The “Viral protein interaction with cytokine and cytokine receptor” pathway was significantly enriched (adjusted p-value < 0.05) in muscle-skeletal.

## Discussion

In this research we present a large-scale study testing the changes in gene expression and exploring marker genes of MV vs. non-MV samples from GTEx [42] donors, across six human tissues—lung, liver, muscle-skeletal, adipose-subcutaneous, skin and nerve-tibial. We developed 18 machine learning models, three models for each of the six tissues, using the XGBoost, RF and NN methods to evaluate their predictive power for MV vs. non-MV samples and for feature analysis purposes. Our results show that the three methods can distinguish MV from non-MV samples successfully for these six tissues and that XGBoost outperforms the other methods, with an average accuracy of 0.951 and average AUC of 0.945 across the six tested tissues. The accuracy and AUC scores for the XGBoost models were higher than the ANN and RF models in all metrics aside from recall, in which the Neural Network model outperforms XGBoost but only by a small margin (all metrics > 0.93). Feature analysis showed that most significant genes affecting the prediction were related to tissue development, movement regulation, fibrosis and inflammation. We furthered explored marker gene convergence across tissues. Further enrichment analysis of the top marker genes showed significant enrichment of cytokine and viral signals for the tested tissues.

The importance of this research is not in the ability to precisely predict the death circumstances based on the gene levels, but rather in the analysis of the features that the machine learning model finds to be significant. Examining the different genes as part of a feature importance analysis reveals unexpected results, in the sense that we detect noteworthy genes or gene combinations with rather distinct value differences between the MV and non-MV groups. We detected tissue specific MV marker genes and marker genes shared across several tissues. Among the shared genes we detected CLEC3B, which is one of the most discriminant genes in both muscle and adipose (see Table 4) and is decreased in the MV group (see

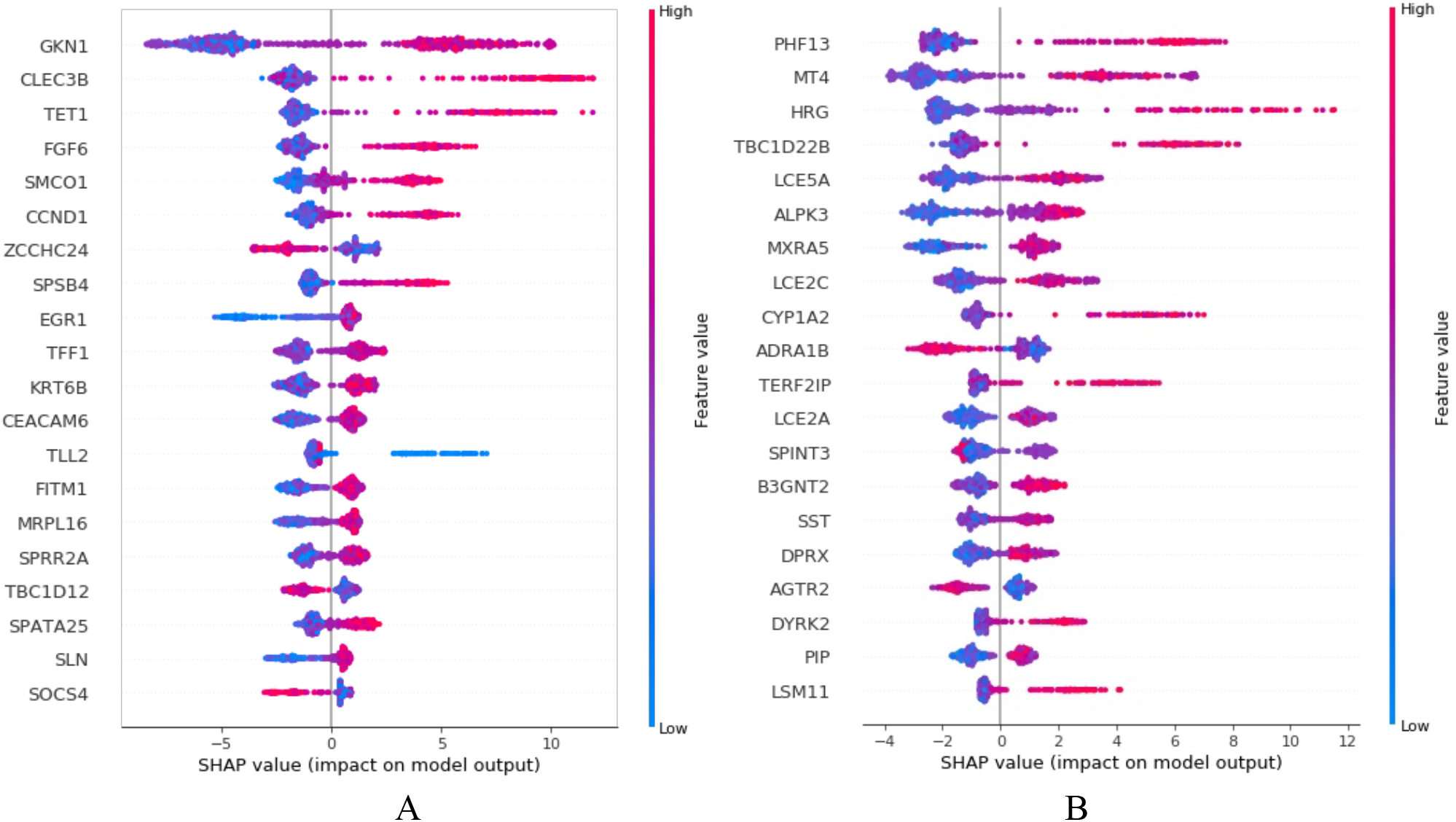

Figure 3A and supplemental Figure S3). The CLEC3B gene, which expresses the Tetranectin protein, has been reported to regulate muscle development [43] and is dysregulated in tumor tissues [45], which may explain its lower values in the MV group (see

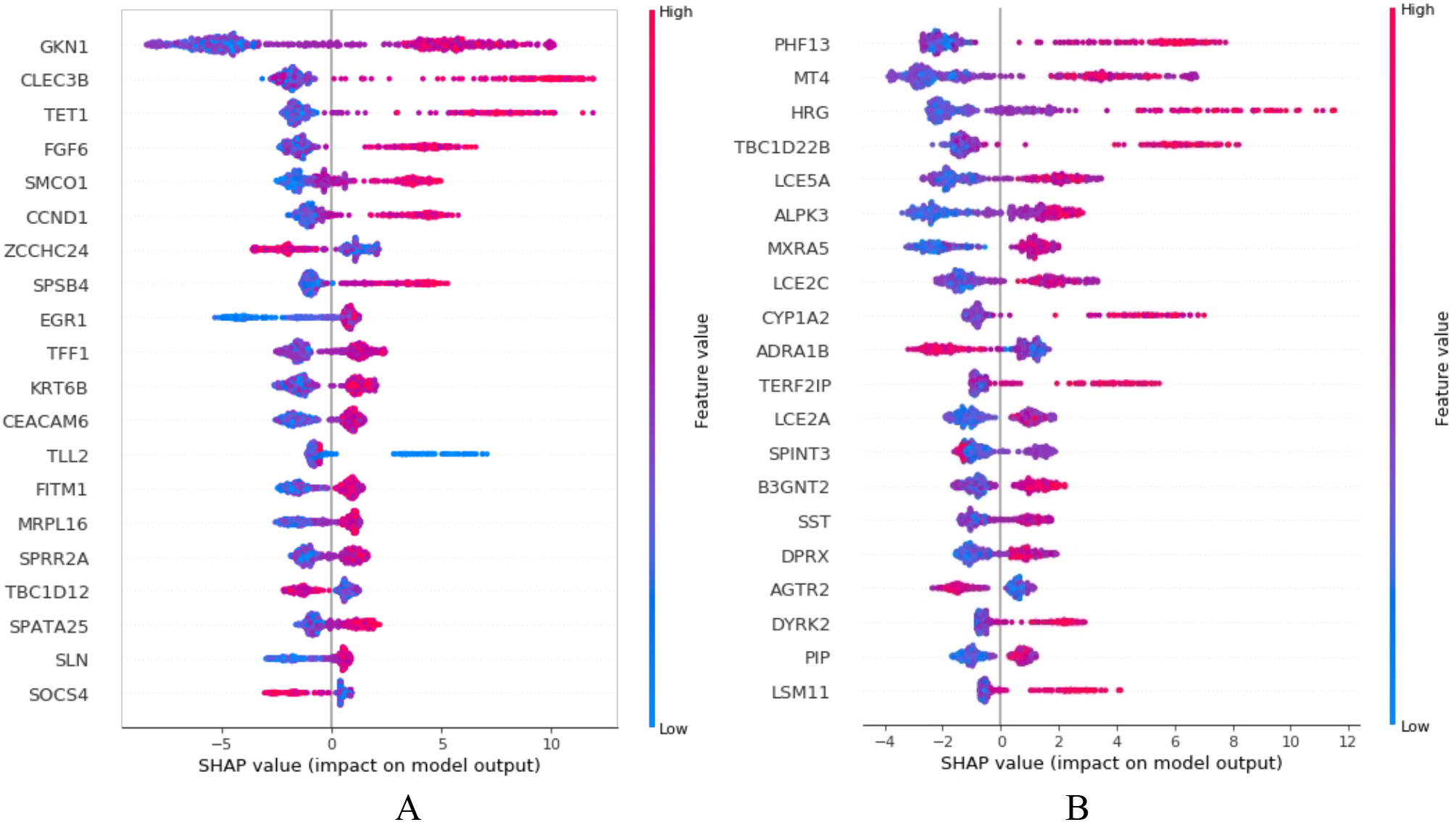

Figure 3A) which is assumed to have decreased muscle development following ventilation. The MXRA5 and EGR1 genes are both strong predictors of ventilation. MXRA5 is a highly significant (see Table 4) gene in the lungs, adipose-subcutaneous and skin-sun exposed tissues and EGR1 in muscle, nerve and adipose. Both the MXRA5 and EGR1 genes were shown to be related to myocardial injury and dysregulated in on-pump vs. off-pump coronary artery bypass surgery [46]. In addition, MXRA5 is an apoptosis and remodeling marker that is associated with the ventilation injury-related response [46] and is a biomarker of severe respiratory syncytial virus (RSV) infection [47]. The tissue-specific angiotensin II receptor 2 (AGTR2) gene is expressed in lung fibrosis [44] and is shown by our analysis to be highly regulated in the lung’s ventilation group (see

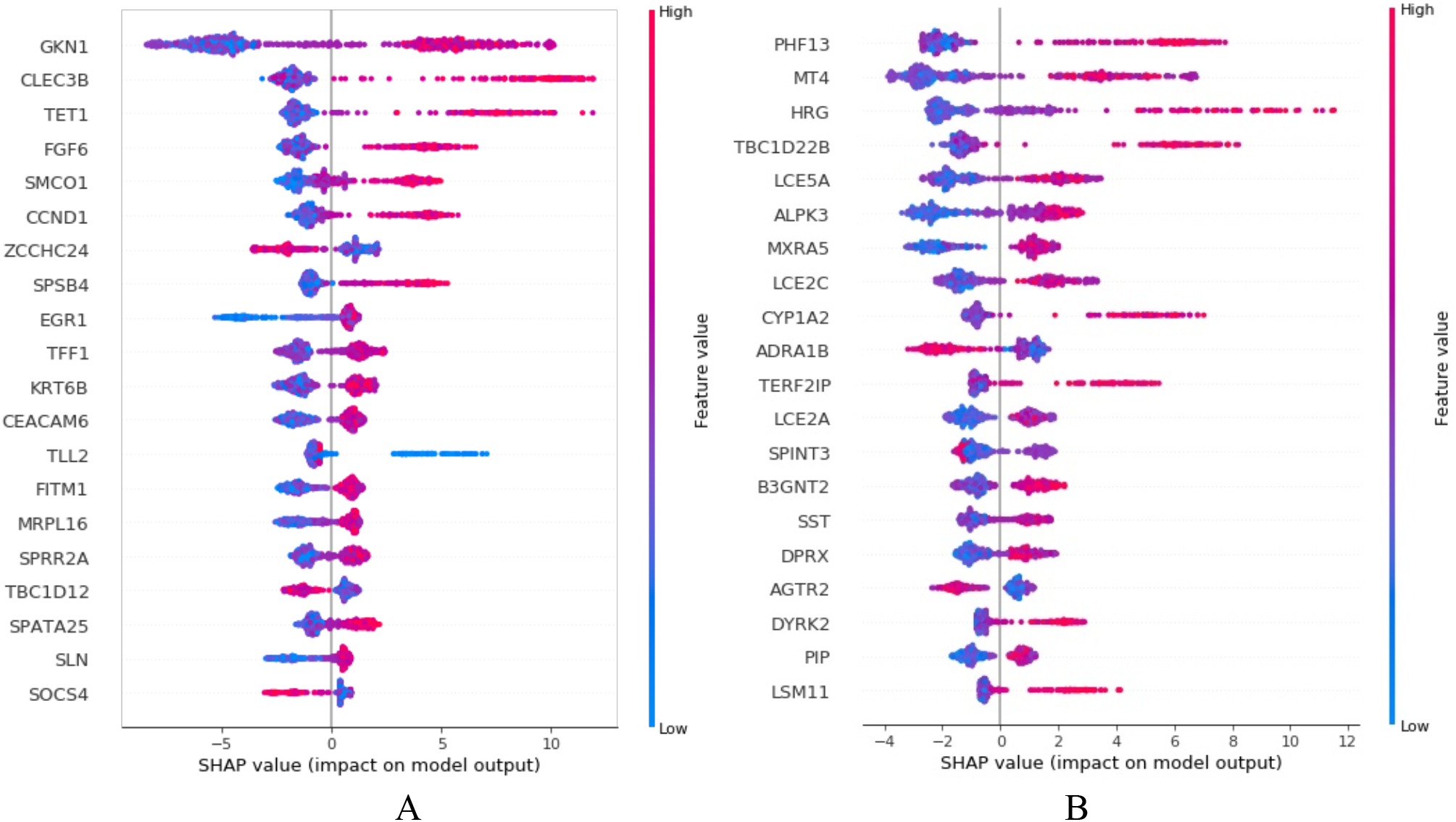

Figure 3B). HRG is a high predictor of MV and is upregulated in the non-MV group (see

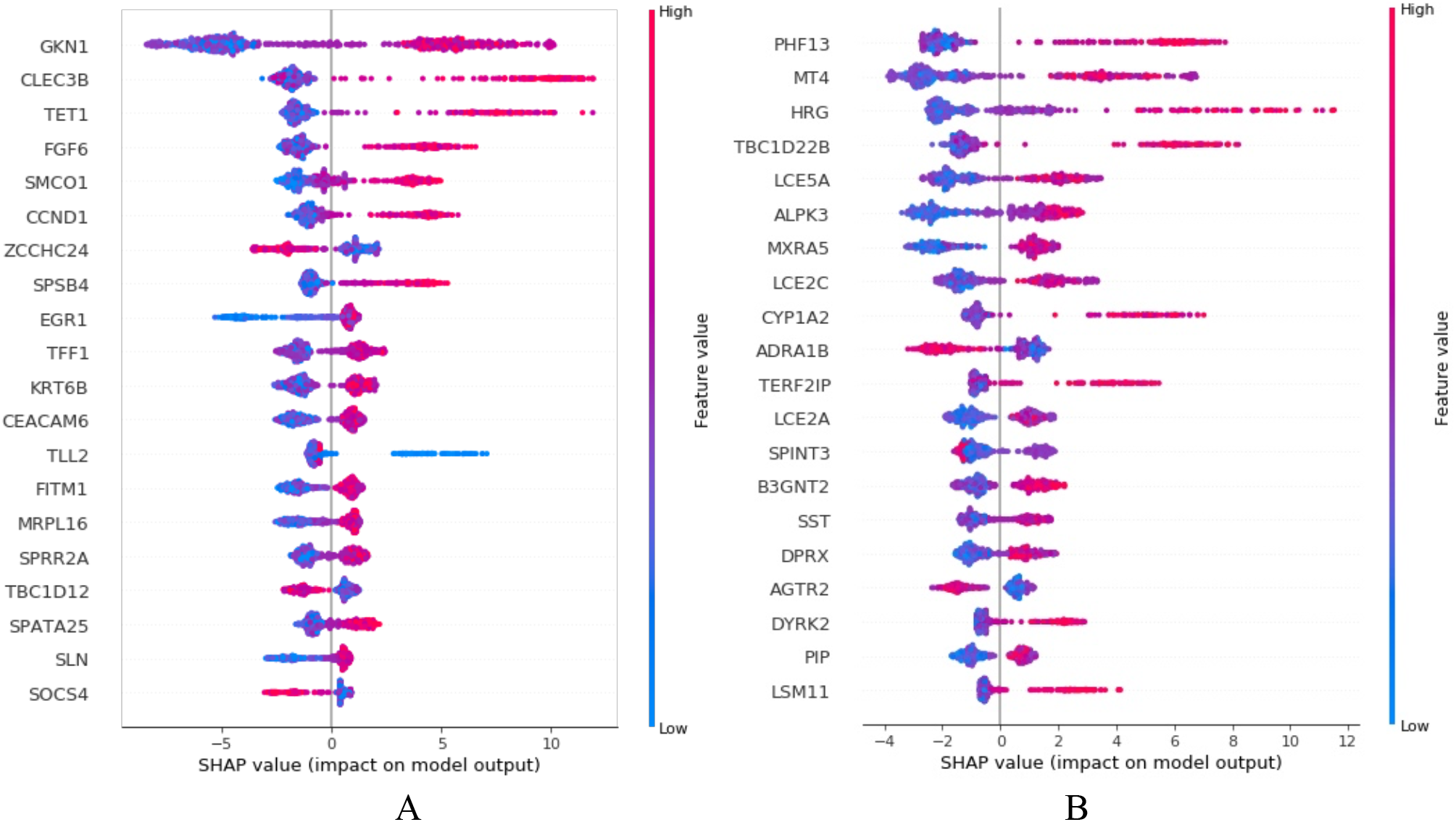

Figure 3B). HRG gene levels are decreased in advanced lung cancer and the gene is known to have antifibrinolytic properties [48] that support its detected low levels in MV samples, which may indicate the progression of lung fibrosis. The TET1 gene, a significant predictor of MV in the muscle, is related to osteogenesis and adipogenesis inhibition [49]. The two top ranked liver marker genes are the TOR1A gene, which is related to smooth physical movements in the brain [49], and DNAJB4, a tumor suppressor gene [50]. In addition, we detected multiple carcinogenic and cancer marker genes among the top high predictor genes of ventilation such as VENTX [51], detected in adipose, MGMT [52] and DNAJB4 [50], detected in the liver, and GKN1 [53], which is an anti-inflammatory protein [54], detected in muscle and adipose with lower expression levels for the ventilation group. Further enrichment analysis of the top genes in each tissue indicated an inflammatory and viral gene signature. In support of our results for the lungs, we mention McCall et al. [13] who, using the lung GTEx gene expression data, detected a large cluster of genes associated with type II pneumocytes related to cells that proliferate in ventilator associated lung injury. One explanation for the inflammatory signature we observed in our findings may be that ventilator-induced lung injury initiates non-pulmonary whole-body organ dysfunction [55].

As research limitations, we detected changes in gene expression between donors that were connected to MV prior to death and donors that were not. We note that these changes may be related to any direct and indirect factor related to the MV constellation, the MV machines, the patient’s extended period of immobility (we indeed detected decreases in gene levels related to movement and development), and any other factor related to being under MV. In addition, since the GTEX tissues samples are derived from post-mortem but healthy donors at time of death, we assume a gene distribution similarity between the deceased and living MV patients.

It is also worth noting that the GTEx tissue data includes bulk gene expression data composed of various cell types. Some gene expression changes we detected may be related to changes in the proportions in the cellular composition of the tissues. For example, the inflammatory signals may be related to changes in the proportions of immune-related cells in the tissues and not merely changes in the expression of genes within the cells. Nevertheless, these marker gene levels predicted MV and non-MV samples successfully and are a significant explanatory tool to understand ventilation induced changes either via gene expression changes within cells or proportion changes of cells in the tissues. We note that we may further improve our performance by utilizing various extensions. For example, Reddy et al. [56] tested four popular machine learning methods and showed that if the dataset is of a high dimensionality, by performing a Principal Component Analysis (PCA) as a preprocessing step, it is possible to reach higher accuracy rates for the model. Nevertheless, our focus is to detect a signature of single genes and not merely increase the performance. Reddy et al. also suggested the Antlion re-sampling-based deep neural network model for classification of an imbalanced multimodal stroke dataset [57]. ElStream [56] is a new approach to detect concept drift utilizing ensemble methods. As further future work, we may use differential network analysis approach to gain further knowledge of the genes networks that changed between the MV and non-MV samples, e.g. Basha et al. [20] used differential network analysis to detect the changes in gene networks across human tissues.

In conclusion, we showed that MV induced transcriptomic changes in six tissues, going beyond the known direct effect on the lungs [21], and related to inflammation, fibrosis, tissue development, growth and movement regulation across all tested tissues. The changes in gene levels that we detected are highly significant and consistent across direct and peripheral tissues and thus MV should be carefully considered before being given to patients.

## Acknowledgements

We thank Dr. Einat Minkov and Dr. Itay Dattner for their consultation. This work was supported by a grant from the DSRC (Data Science Research Center) at the University of Haifa and by an internal fund from the University of Haifa.

## Author contributions

JS and NL designed the experiment. GAY and ES pre-processed and corrected the data. NL conducted the analysis. JS and ES conducted the literature review. JS, NL and ES wrote the paper.

## Data availability

The GTEx data is available for download from (https://www.gtexportal.org/home/datasets).

## Footnotes

1 https://scikit-learn.org/

2 https://keras.io/

3 https://www.tensorflow.org/

4 https://scikit-learn.org/

